# The Absence of Universally-Conserved Protein-coding Genes

**DOI:** 10.1101/842633

**Authors:** Change Laura Tan

## Abstract

Public access to thousands of completely sequenced and annotated genomes provides a great opportunity to address the relationships of different organisms, at the molecular level and on a genome-wide scale. Via comparing the phylogenetic profiles of all protein-coding genes in 317 model species described in the OrthoInspector3.0 database, we found that approximately 29.8% of the total protein-coding genes were orphan genes (genes unique to a specific species) while < 0.01% were universal genes (genes with homologs in each of the 317 species analyzed). When weighted by potential birth event, the orphan genes comprised 82% of the total, while the universal genes accounted for less than 0.00008%. Strikingly, as the analyzed genomes increased, the sum total of universal and nearly-universal genes plateaued while that of orphan and nearly-orphan genes grew continuously. When the compared species increased to the inclusion of 3863 bacteria, 711 eukaryotes, and 179 archaea, not one of the universal genes remained. The results speak to a previously unappreciated degree of genetic biodiversity, which we propose to quantify using the birth-event-weighted gene count method.

## Introduction

The rapid advances of whole genome sequencing technologies have facilitated comparative and evolutionary analyses, at fine molecular detail, mostly based on comparisons of similarities of DNA, RNA, or protein sequences, e.g., homologous genes or gene contents.

Many methods have been developed to identify homologous genes. By definition, any gene and its homologs found in other species are derived from a common ancestral gene. Since the history of genes are generally not known, homology-identification is challenging. Consequently, different methods that are based on different assumptions and algorithms may differ in homology classifications. OrthoInspector is one of the three most balanced methods (the other two are InParanoid and Hieranoid) of orthology inference in specificity and sensitivity (1–3).

OrthoInspector identifies orthologs by dividing genes into inparalog groups based on all-to-all proteome BLAST comparisons and then searching for a reciprocal-best-hit relationship between inparalog groups (1). Since it does not require a reciprocal-best-hit between individual genes it is more sensitive than many other methods, including Inparanoid and OrthoMCL (1, 2, 4, 5). Recently, the OrthoInspector algorithm has been used to determine orthologs in 4753 organisms (3863 bacteria, 711 eukaryotes, and 179 archaea), generating an orthology resource with the broadest species coverage (except viruses) (6). Of the 4753 organisms, 317 (144 eukaryotes, 142 bacteria, and 31 archaea) are deemed model species (referred to as NMS (Nevers’s Model Species) hereafter) either due to their importance in the biological field or due to a consideration of taxonomic coverage. Orthologous genes across three domains of life are available in the OrthoInspector website for these species. For a non-model species, only orthologous genes within its own domain of life are available.

To investigate the relationship of gene contents of different species, we compared the phylogenetic profiles of all NMS. A phylogenetic profile of a protein describes the presence or absence of its homologs across a given set of organisms (7). Two proteins with the same phylogenetic profiles tend to function in the same biological process, though the accuracy of the functional-linkage prediction depends on the criteria of defining homology and selection of reference species (7–11). We focused our attention to proteins with two extreme distribution: universal or orphan. To accommodate the potential bias of the method used to identify homologous genes and the consequence of occasional gene loss or genome sequencing or annotation errors, we have also analyzed nearly-universal genes and nearly-orphan genes.

We discovered an unexpected pattern of the distribution of universal genes and orphan genes. We found that every species has a large number of orphan and nearly-orphan genes, but none, or only a few, universal and nearly-universal genes. Contrary to the common expectation that homologs would be found for orphan genes so that orphan gene number would decrease as more species are analyzed, the number of orphan genes grows continuously; each addition of species brings new orphan genes, though often resulting in a decrease of universal genes. Strikingly, not only the homologs of a universal gene are generally not universal genes, but also not a single universal gene maintained its status as a universal gene when enough species were sampled. In other words, all genes are taxonomically restricted, though at different levels of restriction.

## Materials and Methods

Phylogenetic profiles of all the 317 NMS were provided by Nevers and Lecompte in CSV (delimited by tabs). The phylogenetic profile for each species contains information about the presence (indicated with a 1) or absence (indicated with a 0) of homologs in the 317 NMS (columns) for all its protein-coding genes (rows). For the column that corresponds to the species itself, all cells are 0. When the CSV files were imported into Excel, all species names were shifted one cell to the left. After that was corrected, the phylogenetic profiles were saved as Excel files. Number of species in which there are orthologous genes for a NMS protein was calculated based on the phylogenetic profiles using Microsoft Excel and/or a script written for this project by Andrew Jones. Identity of proteins and that of their orthologs were manually curated from the OrthoInspector website (https://lbgi.fr/orthoinspectorv3/). Gene function annotations were mostly from the UniProt Knowledgebase (UniProtKB, https://www.uniprot.org/uniprot/), occasionally from organism-specific databases, e.g., the Drosophila Genome Database (https://flybase.org/) and the Saccharomyces Genome Database (https://www.yeastgenome.org/). All figures were generated using Microsoft Excel and PowerPoint.

### Categorizing Genes according to the Numbers of Species Having Their Homologs

The number of species having homologs for a specific gene is the sum total for the row to which that gene belongs, since the presence or absence of its homologs in a species (except its own host or home species) is indicated with 1 or 0, respectively. Note that the presence of multiple homologs in a species does not increase the count beyond 1. A gene was called an orphan, orphan +1, orphan +2…or a universal (o+1, o+2…universal, “ o” means orphan) gene if the sum total is 0, 1,2. or 316.

### Identification of Orphan, Nearly-orphan, Universal, and Nearly-universal Genes

If the number of species having homologs for a specific gene is 316, then that gene is a universal gene, since it has homolog(s) in every organism analyzed. If the number is 0, then that gene is an orphan gene, a gene unique to a species; no homolog exists in any of the other species analyzed. Nearly-universal genes are genes conserved in all but five or fewer species analyzed, i.e., a sum total of 315, 314, 313, 312, or 311 (corresponding to u-1, u-2, u-3, u-4, u-5 genes, “ u” means universal). Nearly-orphan genes, by contrast, are genes that are shared by no more than five of the species analyzed, i.e., a sum total of 1,2, 3, 4, or 5 (corresponding to o+1, o+2, o+3, o+4, o+5 genes).

### Weighted Counts of Genes by Potential Birth Event

The weighted value of a gene is the inverse of the number of species, including the gene’s home species, that have homologs for that gene. Thus, an orphan gene was counted as one (=1/1) gene, while a universal gene was counted as 0.003155 (=1/317) gene. An o+x gene was counted as 1/(1+x) gene, x is any integer between 1 and 316.

## Results

### Distribution of Total Genes

To gain a broad view of the species being analyzed, we compared the sizes of their proteomes (Fig 1). Not surprisingly, on average, eukaryotes have much larger proteome sizes than bacteria and archaea. Eukaryote proteome sizes also vary the most.

**Fig 1.**
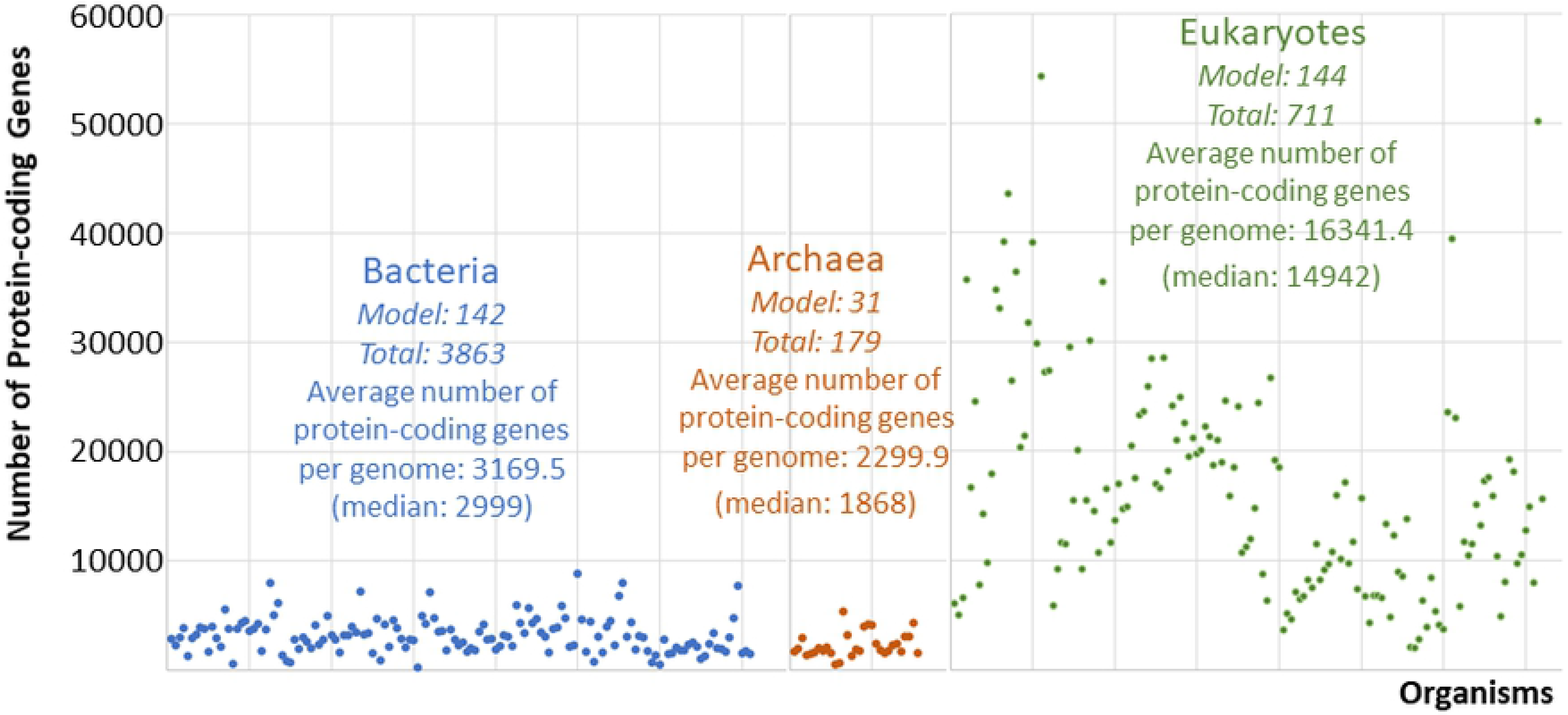
Number of protein-coding genes of NMS. Each data point represents a species. Species are arranged according to the order of (6). The first two numbers underneath the name of each domain of life are the number of the model species and that of the total species analyzed in the corresponding domain.

### Grouping Genes according to the Occurrences of Their Homologs

Next, we placed every protein-coding gene encoded in the genomes of all NMS into orphan, orphan +1, orphan +2… or universal groups (o, o+1, o+2…universal) based on whether the number of species in which its homologs exist is 0, 1,2… or 316 (S2 Table). Surprisingly, for every species analyzed, the group with most members was the orphan group, and the number of genes in a group quickly dropped into a handful or even zero with the increase of species containing the corresponding homologs (Fig 2 and S1 Fig).

**Fig 2.**
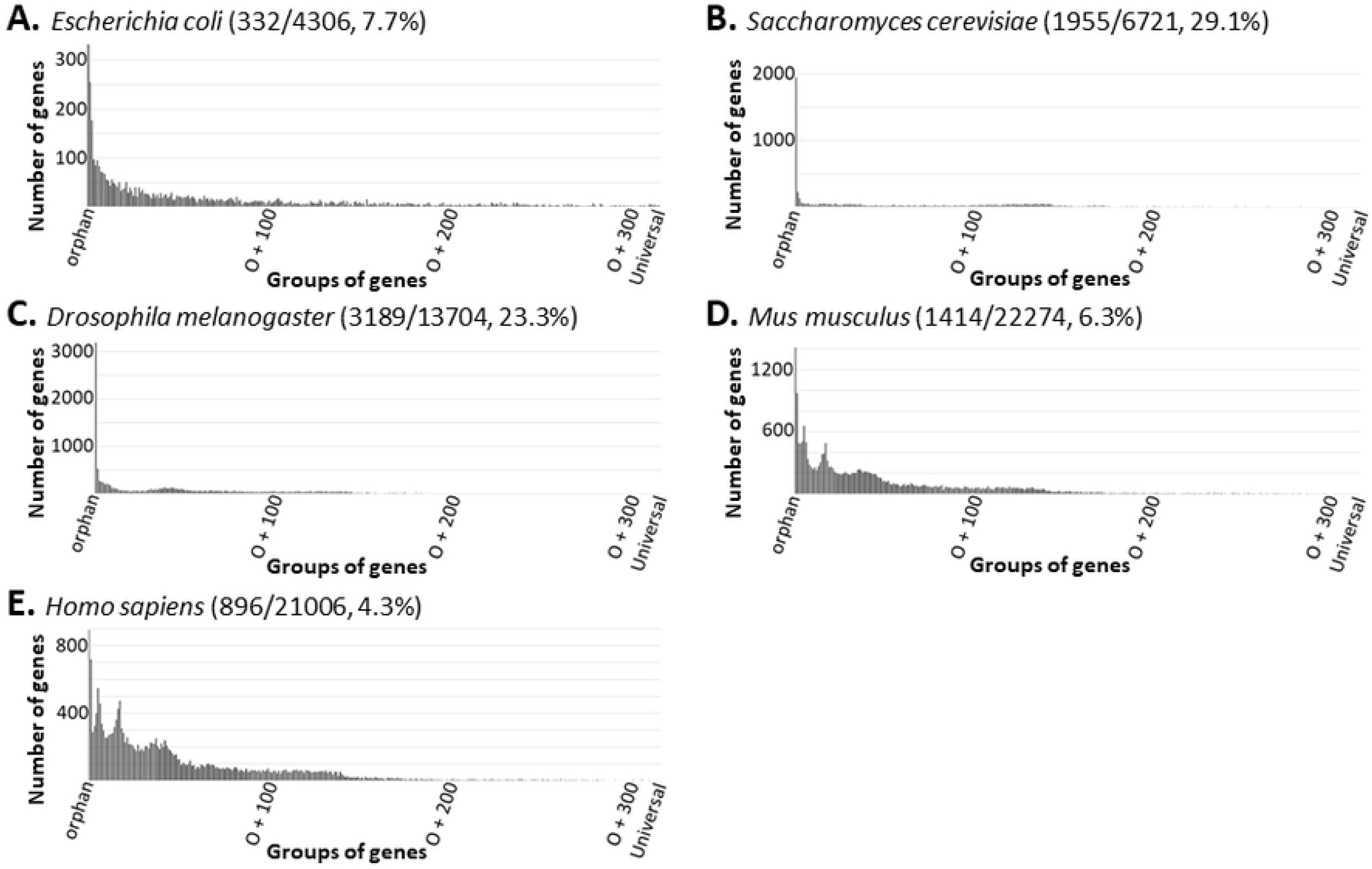
The number of genes in a group decreases rapidly as the number of species sharing the corresponding homologs increases. Note that the first vertical line in each panel is not its Y-axis but the orphan gene number in the corresponding organism. The numbers in the parenthesis next to the name of a species in each panel are the number of its orphan genes, all protein-coding genes, and the percent of orphan genes. Only five species are shown here. More examples can be found in S1 Fig and S2 Table.

### Distribution of Universal, Nearly-universal, Orphan, and Nearly-orphan Genes

In order to comprehend the above data, we focused our attention on orphan, nearly-orphan, universal, and nearly-universal genes. A nearly-universal gene is conserved in all but five or fewer species analyzed, while a nearly-orphan gene is shared by no more than five of the species analyzed. As one would expect from a quick glance at Fig 2, S1 Fig, and S2 Table, the number of orphan and nearly-orphan (ONO) genes greatly exceeded that of universal and nearly-universal (UNU) genes. Table 1 lists the ONO, UNU, and the proteome of our seventeen chosen organisms, including three bacteria, three archaea, three plants, one fungus, and seven animals. S3 Table lists the ONO, UNU, and the proteome of all NMS. Fig 3 shows the percentages of ONO genes (colored sections at the bottom), UNU genes (colored sections at the top), and all other genes (gray) in each of the model species. Note the great portion of the ONO genes. In contrast, the portion of the UNU genes are barely visible, especially for eukaryotes.

**Fig 3.**
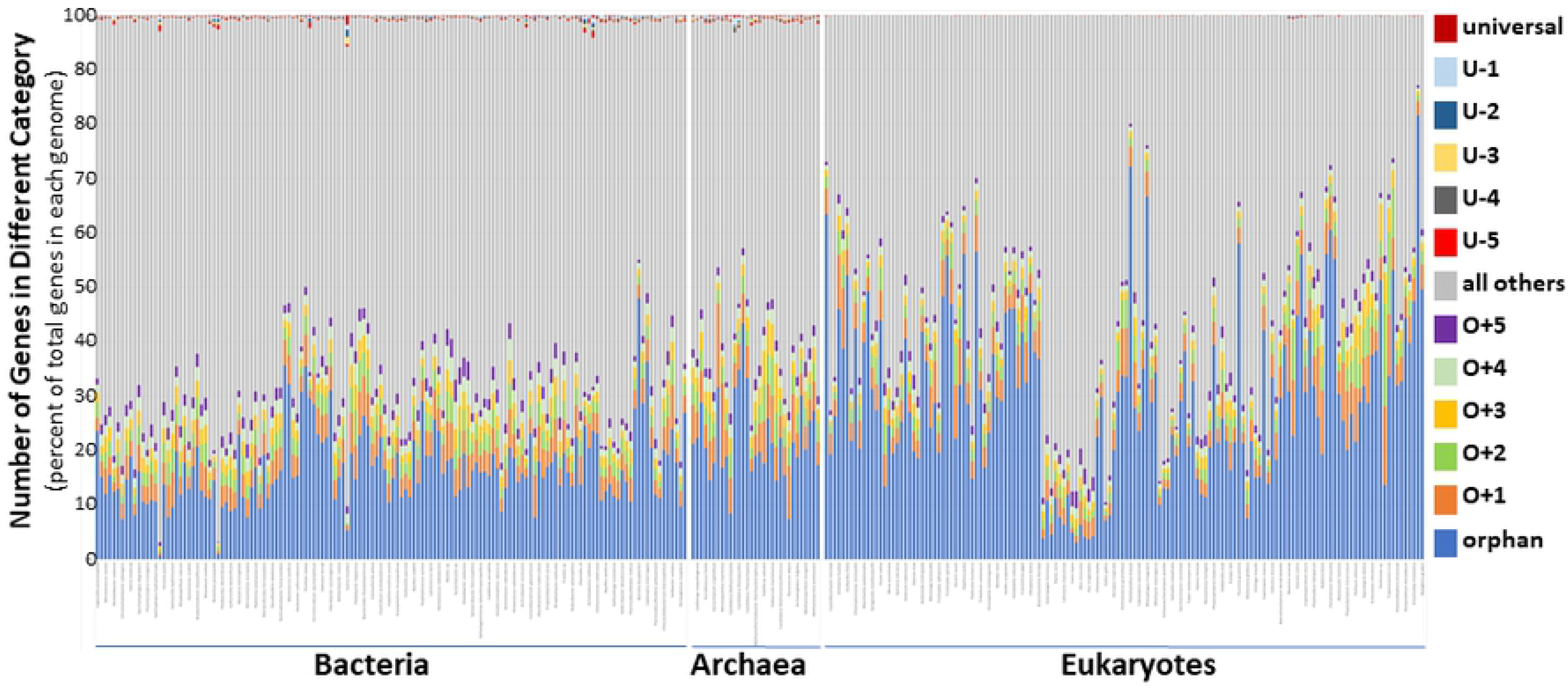
Distribution of ONO and UNU genes of all NMS. Each column represents a species whose order of appearance is according to that of (6). Not all organisms are labeled and the species names are too small to read in this figure due to the limitation of space. For a higher resolution figure with all organisms legibly labeled, see the big graph located at the bottom of S3 Table.

**Table 1:** Numbers of orphan, nearly-orphan, universal, and nearly-universal genes in seventeen chosen organisms

| Species | o | o+1 | o+2 | o+3 | o+4 | o+5 | u-5 | u-4 | u-3 | u-2 | u-1 | u | Proteome |
| --- | --- | --- | --- | --- | --- | --- | --- | --- | --- | --- | --- | --- | --- |
| *Escherichia coli* | 332 | 255 | 177 | 97 | 85 | 95 | 4 | 3 | 4 | 2 | 3 | 1 | 4306 |
| *Caulobacter crescentus* | 635 | 130 | 99 | 86 | 81 | 74 | 5 | 1 | 2 | 0 | 6 | 0 | 3715 |
| *Mycoplasma genitalium* | 114 | 16 | 11 | 4 | 5 | 3 | 5 | 2 | 2 | 4 | 5 | 2 | 483 |
| *Sulfolobus solfataricus* | 843 | 169 | 108 | 102 | 69 | 51 | 3 | 3 | 2 | 2 | 4 | 1 | 2924 |
| *Methanocaldococcus jannaschii* | 353 | 94 | 60 | 59 | 36 | 34 | 4 | 4 | 4 | 5 | 3 | 4 | 1787 |
| *Haloferax volcanii* | 700 | 311 | 265 | 248 | 116 | 60 | 5 | 4 | 3 | 2 | 4 | 4 | 3982 |
| *Zea mays* | 10719 | 1869 | 1933 | 1033 | 863 | 761 | 3 | 0 | 0 | 0 | 0 | 0 | 39154 |
| *Oryza sativa* | 19132 | 2380 | 1847 | 958 | 740 | 627 | 2 | 2 | 0 | 0 | 0 | 0 | 43550 |
| *Arabidopsis thaliana* | 5057 | 893 | 558 | 544 | 546 | 545 | 0 | 1 | 0 | 0 | 0 | 0 | 27252 |
| *Saccharomyces cerevisiae* | 1955 | 225 | 128 | 61 | 45 | 44 | 3 | 2 | 0 | 0 | 1 | 0 | 6721 |
| *Caenorhabditis elegans* | 9418 | 1285 | 994 | 355 | 201 | 180 | 2 | 1 | 0 | 0 | 1 | 0 | 20071 |
| *Drosophila melanogaster* | 3189 | 525 | 269 | 250 | 238 | 195 | 1 | 4 | 0 | 0 | 0 | 0 | 13704 |
| *Danio rerio* | 2808 | 744 | 496 | 488 | 449 | 339 | 0 | 2 | 0 | 0 | 0 | 0 | 24929 |
| *Canis lupus* | 973 | 171 | 188 | 283 | 311 | 534 | 2 | 1 | 0 | 0 | 0 | 0 | 19756 |
| *Mus musculus* | 1414 | 972 | 489 | 488 | 510 | 658 | 2 | 2 | 0 | 0 | 0 | 0 | 22274 |
| *Homo sapiens* | 896 | 719 | 288 | 325 | 402 | 548 | 1 | 2 | 0 | 0 | 0 | 0 | 21006 |
| *Gallus gallus* | 1136 | 143 | 150 | 79 | 87 | 77 | 1 | 2 | 0 | 1 | 1 | 0 | 15913 |

Fig 4 is a representation of the grouping of genes of all NMS together. Amazingly, the ONO genes represent 42.7% (1,228,529) of the total (2,874,537), with the orphan group itself occupies about 29.8% (855,723) of the total. The UNU groups account for less than 0.14% (3,906) of the total, with the universal group about 0.01% (261) of the total.

**Fig 4.**
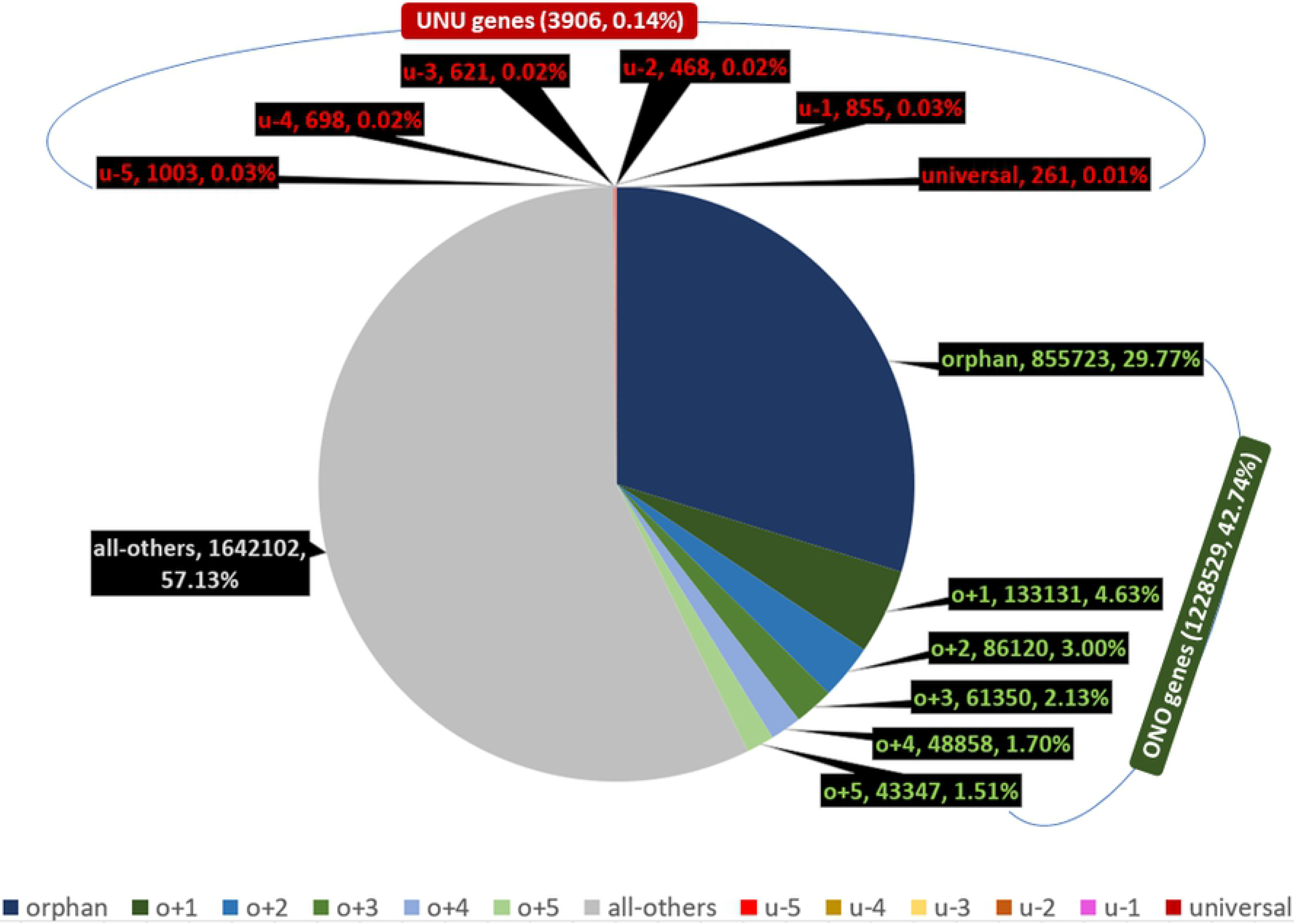
Distribution of orphan, nearly-orphan, universal, and nearly-universal genes of NMS as a whole.

Strangely, the number of universal genes, 261, is even smaller than the number of model species, 317. This creates a contradiction with the definition of universal genes, since, by definition, if one true universal gene existed in the model species, then we should have 317 universal genes, because a universal gene should (again, by definition) possess a homolog in each of the other model species. To make the situation worse, more than half of the model species have none, while some of them have several universal genes (Table 2). For example, *Candidatus caldiarchaeum* has seven universal genes, almost twice as many as the ten species with the second largest number (four) of universal genes. Although eukaryotic proteomes are generally much larger than bacterial and archaeal proteomes, they have the least number of universal genes. Of the 144 model eukaryotes, only ten have universal genes and none has more than one universal genes.

**Table 2:** Distribution of species with zero to seven universal genes

| Domain of Life |  | Bacteria | Archaea | Eukaryotes | All |
| --- | --- | --- | --- | --- | --- |
| Number of Model Species |  | 142 | 31 | 144 | 317 |
| Number of Universal Genes | 0 | 33 | 1 | 134 | 168 |
|  | 1 | 59 | 4 | 10 | 73 |
|  | 2 | 45 | 9 | 0 | 54 |
|  | 3 | 3 | 8 | 0 | 11 |
|  | 4 | 2 | 8 | 0 | 10 |
|  | 5 | 0 | 0 | 0 | 0 |
|  | 6 | 0 | 0 | 0 | 0 |
|  | 7 | 0 | 1 | 0 | 1 |
| Universal Genes per Genome |  | 1.17 | 2.74 | 0.07 | 0.82 |

### Weighted Distribution of UNU and ONO Genes

Homologs of a gene are generally thought to have shared a common gene ancestor. In other words, they shared one “ birth event”. Therefore, it is only logical that genes should not have been counted equally; they should be weighted according to their birth events. Consequently, each orphan gene shares its birth right with no other sequence, and should be counted once, while each o+1 gene shares its birth right with an ortholog, and should be counted as 0.5, and so on. When thus weighted, the ONO genes represent about 95% of the total, while the UNU groups represent only about 0.0012% (Fig 5 and S4 Table). The orphan group itself makes up more than 82% of the total. In comparison, without weighting, when every gene is counted equally in each species, the ONO genes represent about 43% of the total, while the UNU genes account for about 0.14%.

**Fig 5.**
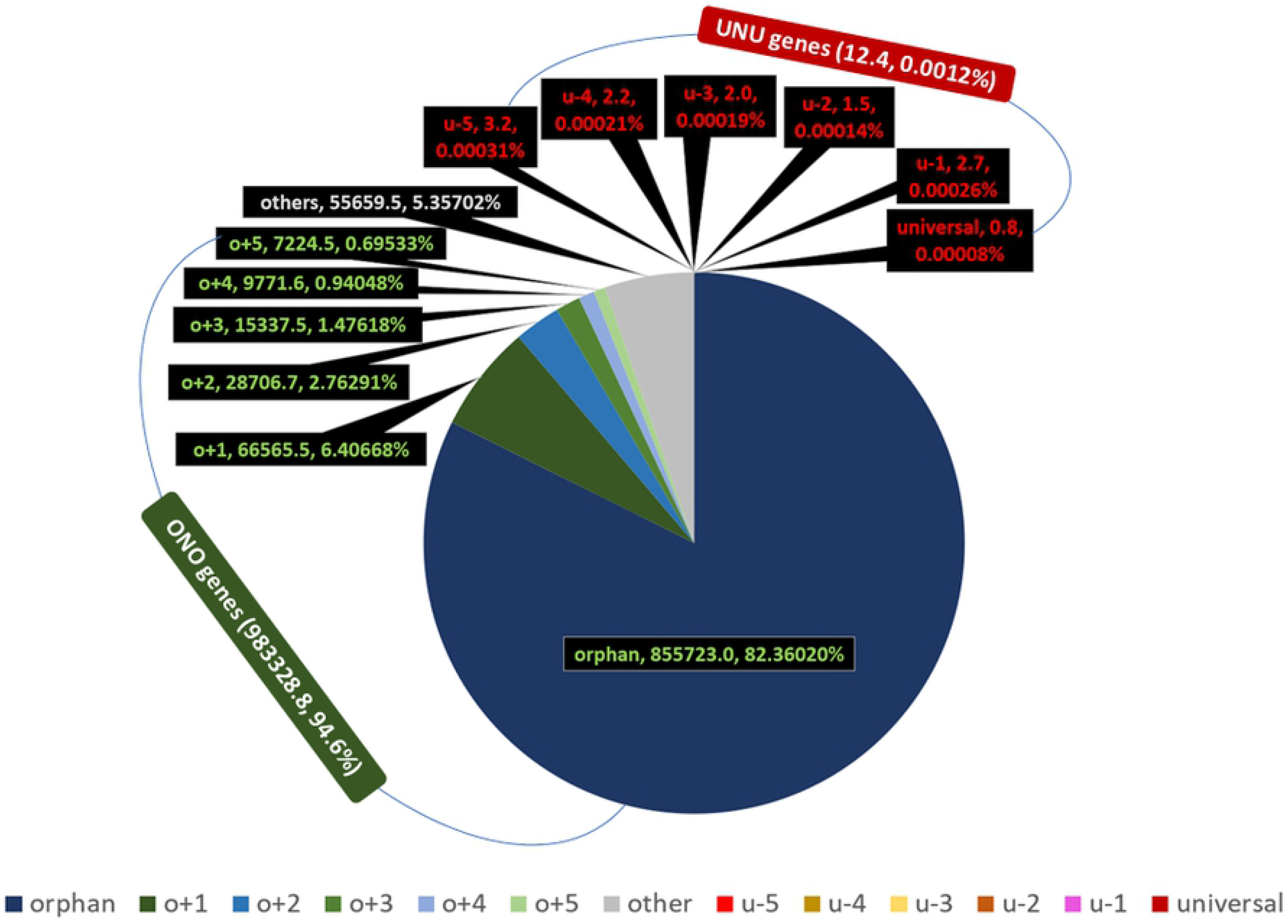
Distribution of weighted orphan, nearly-orphan, universal, and nearly-universal genes of NMS as a whole.

### Change of the Total Numbers of UNU and ONO Genes with the Species Coverage

To determine the accumulated number of UNU, ONO, and all the protein-coding genes as more species were added, we simply summed the UNU, the ONO, and the proteome of each of the model species, one-by-one, in the sequence of the Nevers’s original species order. The number of UNU grew quickly initially with the addition of species, but the growth slowed down soon, and almost plateaued at around 3,300 genes (Fig 6A). Strikingly, the ONO number increased continuously, at a much greater speed than the initial, fastest, growth rate of the UNU number (Fig 6B, orange). Viewed with the same scale, the number of UNU genes appears to show a trend, or slope, of zero (i.e., unchanging along the vertical axis) (Fig 6B, blue).

**Fig 6.**
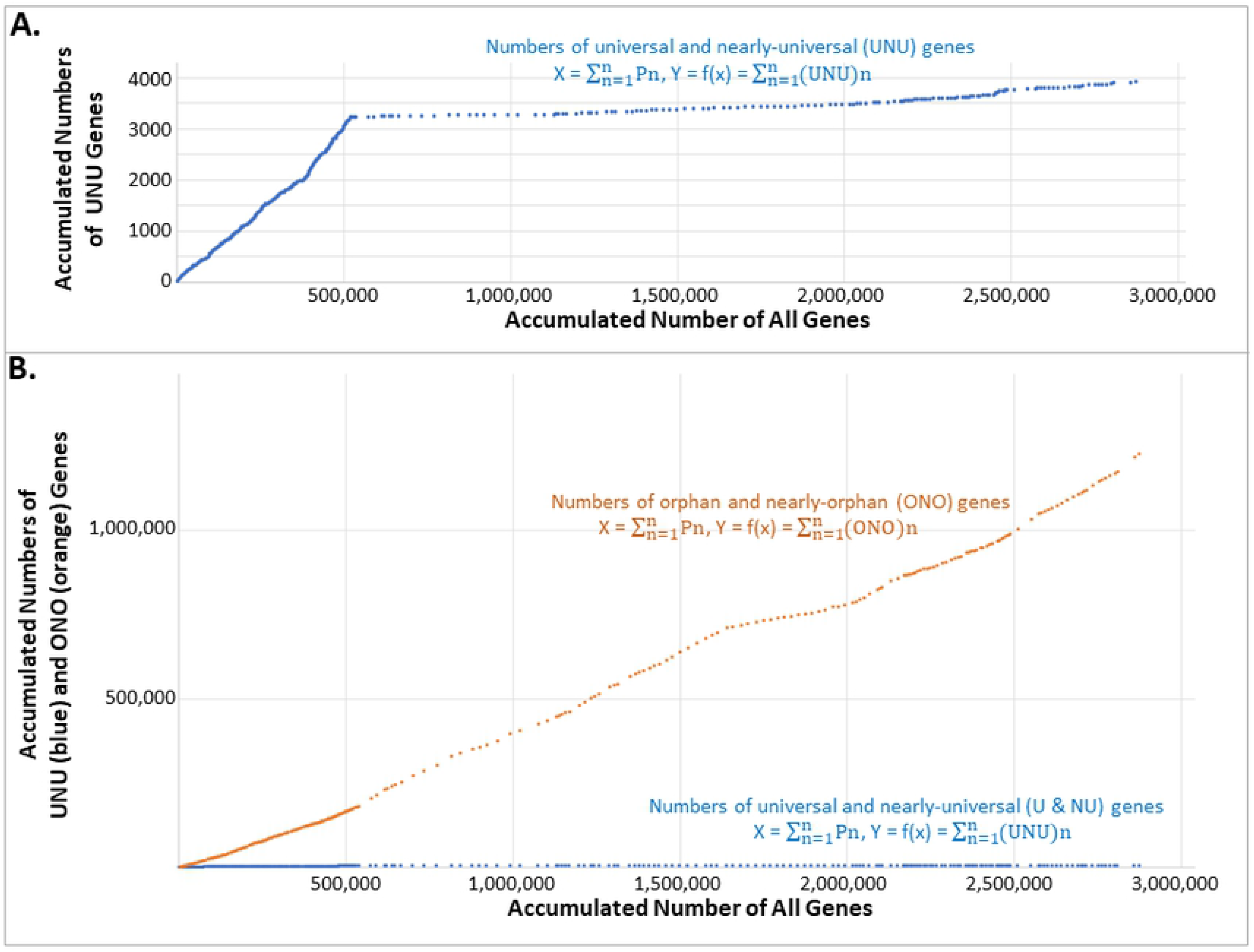
Change of the number of UNU, ONO, and all the protein-coding genes as new organisms were added. Each data point represents a species whose coordinates are (X, Y) and whose encoded protein number is Pn. X is the sum of the proteins encoded by all the organisms up to and including that species. Y is the sum of the ONO or UNU of these organisms.

### Identity of the Universal Genes

The numbers of different groups of genes are interesting and important to know but their identity is even more informative for our understanding of life. Here we will describe the identity of the universal genes, while that of the orphan genes will be described in another publication.

We first analyzed the homologs of the universal genes of our chosen 17. We observed that the vast majority of the homologs of a gene encode proteins perform the same function in different organisms, when functional data are available (S5 Table). This indicates the specificity of homolog inference of OrthoInspector. The occasional out-of-place homologs, e.g., the *Escherichia coli* lysine-tRNA ligase amongst the homologs of asparagine-tRNA ligase and the *Caulobacter crescentus* peptide chain release factor 3 in the mist of elongation factor G homologs, indicates the ability of OrthoInspector to recognize even very low similarity. For example, the *E. coli* lysine-tRNA ligase, though not an asparagine-tRNA ligase, belongs to the same subgroup (class IIb) of tRNA ligases as the asparagine-tRNA ligase, with the sole other member of this group being aspartate-tRNA ligase (12). These three tRNA ligases recognize related anticodons: aspartate GUC, asparagine GUU, and lysine UUU triplets. The *C. crescentus* peptide chain release factor 3 belongs to the same subfamily of GTPases and probably has similar structure and binds to overlapping regions of the bacterial ribosomes as the elongation factor G (13–15). The fact that most homologs of a gene encode proteins with the same characteristics and/or functions demonstrates that OrthoInspector is a reliable method to identify homologs.

The second phenomenon we observed is that the homolog of a universal gene is commonly not a universal gene. This is surprising. However, it explains why the number of universal genes is not an integer multiplication of the number of model species (317), though not why it is smaller than 317.

The third observation is that none of the universal genes in our chosen 17 kept their status as universal genes when checked against their in-domain non-model species in OrthoInspector (S5 Table).

Next, we expanded our analysis to all the model species. Consistent with the observation of the chosen 17, all universal genes lost their status as a universal gene when checked against their in-domain non-model species in OrthoInspector.

The protein characteristics of five universal genes (Q74MY3_NANEQ, R1E424_9ARCH, E4WXB9_OIKDI, C4V6P4_NOSCE, A2ER26_TRIVA) were not clear. To solve this problem, we examined their homologs in the chosen 17 (S6 Table). Assuming a gene’s homologs share its identity, we called Q74MY3_NANEQ and R1E424_9ARCH Obg-like ATPase 1, A2ER26_TRIVA and C4V6P4_NOSCE elongation factor 2, and E4WXB9_OIKDI isoleucine-tRNA ligase.

With the new assignation for these five, the 261 universal genes encode eight proteins: aspartate-tRNA ligase, phenylalanine-tRNA ligase alpha subunit, valine-tRNA ligase, isoleucine-tRNA ligase, elongation factor G (name according to bacteria, corresponding to the archaeal and eukaryotic elongation factor 2), elongation factor Tu (name according to bacteria, corresponding to the archaeal and eukaryotic elongation factor 1), DNA-directed RNA polymerase subunit beta, and Obg-like ATPase 1 (Table 3 and S7 Table). Five of these eight belong to the Up-to-date bacterial core gene set (the two elongation factors and valine-tRNA ligase do not) (16).

**Table 3:** Identity of universal genes and their distribution in the three domains of life

| Bacteria | Archaea | Eukaryotes |
| --- | --- | --- |
| Aspartate-tRNA ligase (1) | Aspartate-tRNA(Asp/Asn) ligase (10) |  |
| DNA-directed RNA polymerase subunit beta (3) | DNA-directed RNA polymerase subunit beta (5) | DNA-directed RNA polymerase subunit beta (4) |
| Elongation factor G (Elongation factor 2) (7) | Elongation factor 2 (31) | Elongation factor 2 (5) (A2ER26_TRIVA, C4V6P4_NOSCE) |
| Elongation factor Tu (Elongation factor 1) (57) | Elongation factor 1-alpha (11) |  |
|  | Isoleucine-tRNA ligase (19) | Isoleucine-tRNA ligase (1) (E4WXB9_OIKDI) |
| Phenylalanine-tRNA ligase alpha subunit (97) | Phenylalanine-tRNA ligase alpha subunit (7) |  |
| Valine-tRNA ligase (1) |  |  |
|  | Obg-like ATPase 1 (2) (Q74MY3_NANEQ, R1E424_9ARCH) |  |

*Note: The numbers in parenthesis after the names of genes indicate the numbers of species in which the corresponding genes were recognized as universal genes. Genes recognized as universal genes in species across three domains are shaded in dark gray and those between two domains in gray. The original names for the five universal genes that we assigned new identity have also been included in the table.*

Therefore, even though by definition, if there is one universal gene in the model species, we should have 317 such genes because it should have a homologous gene in each of the model species. And all of the 317 homologous genes should be the same, at least very similar. But we have only 261 universal genes and they are not all the same, although they all, except one (Obg-like ATPase 1), are involved in either gene transcription (the DNA-directed RNA polymerase subunit beta) or gene translation (the t-RNA ligases and elongation factors). Even Obg-like ATPase 1 may be involved in translation. For example, human Obg-like ATPase 1 can prevent eIF2 (eukaryotic initiation factor 2) ternary complex formation, leading to inhibition of protein synthesis and promotion of integrated stress response (17, 18).

## Discussion

Our comparisons of the genetic profiles of 317 proteomes revealed that ONO genes are a common occurrence, while the UNU genes are very rare. The more organisms are included in the analysis, the more ONO genes are detected and the smaller the percentage of UNU gene becomes. Lastly, not a single universal gene remains its status as universal when enough organisms are included.

### Universal vs Non-universal Genes

The continuous increase of the ONO numbers and the leveling off of the UNU numbers is consistent with earlier observations (19–29). These analyses have led to a dramatic shrinking, or even vanishing, of the “ universal,” or universally-conserved, core set of genes and proteins – with a concomitant linear growth in the so-called “ orphan” or “ taxonomically restricted” sequences. “ Accordingly,” notes Koonin, “ the universal core of life has shrunk almost to the point of vanishing” (30). Indeed, after complete sequencing of the first two bacterial genomes, a comparison of the 1727 protein-coding genes of *Haemophilus influenza* and the 468 *Mycoplasma genitalium* genes identified 240 homologous genes between the two (24). When the number of included prokaryotic genomes increased to 100, the number of universally-conserved homologous genes decreased to 63 (25). With the inclusion of 1000 genomes, the number of universally-conserved homologous genes became zero — not a single protein-coding gene was conserved across the 1000 prokaryotes compared (26).

What is surprising, and counterintuitive, is that the homologs of universal genes normally are not universal genes. This results from how we define and detect homologs. Two genes are deemed homologous as long as part of their encoded proteins share some sequence similarity, normally an e-value of 10^−3^ to 10^−5^ in a BLASTp search. OrthoInspector uses an e-value cutoff of 1e^−9^ (about 1.2×10^−4^) (1). At this condition, the *C. crescentus* peptide chain release factor 3 and elongation factor G are detected as homologs because they share some sequence homology in their GTPase domains. With the normal e-value cutoffs, the well-known *Drosophila* orphan genes *Jingwei* and *Zeus* will not be recognized as orphan genes because their high sequence similarity with other wildly distributed genes (31–38). For example, *Drosophila melanogaster Zeus,* though without homologs in the bacterial and archaeal model species, has homologs in 100 of Nevers’s 143 eukaryotic model species, including *Saccharomyces cerevisiae, Caenorhabditis elegans*, and *Gallus gallus* (Tan, unpublished observation).

To illustrate how the homologs of a universal gene can be non-universal genes, opposite to what one would expect by the definition of a universal gene, we made up three hypothetical proteins, A, B, and U (Fig 7). U shares parts 1,2, and 3 with A and parts 1, 4, and 5 with B. Each part can be one or a group of amino acids. Under the criterion that two proteins are homologous if they share three parts, U and A are homologs, so are U and B. However, A and B only share part 1, thus are not homologous. U is a universal gene in this scenario, while A and B are not. How would the ancestor(s) of A, B, and U look like, the one on the bottom left, the one on the bottom right, or something else? How can we know?

**Fig 7.**
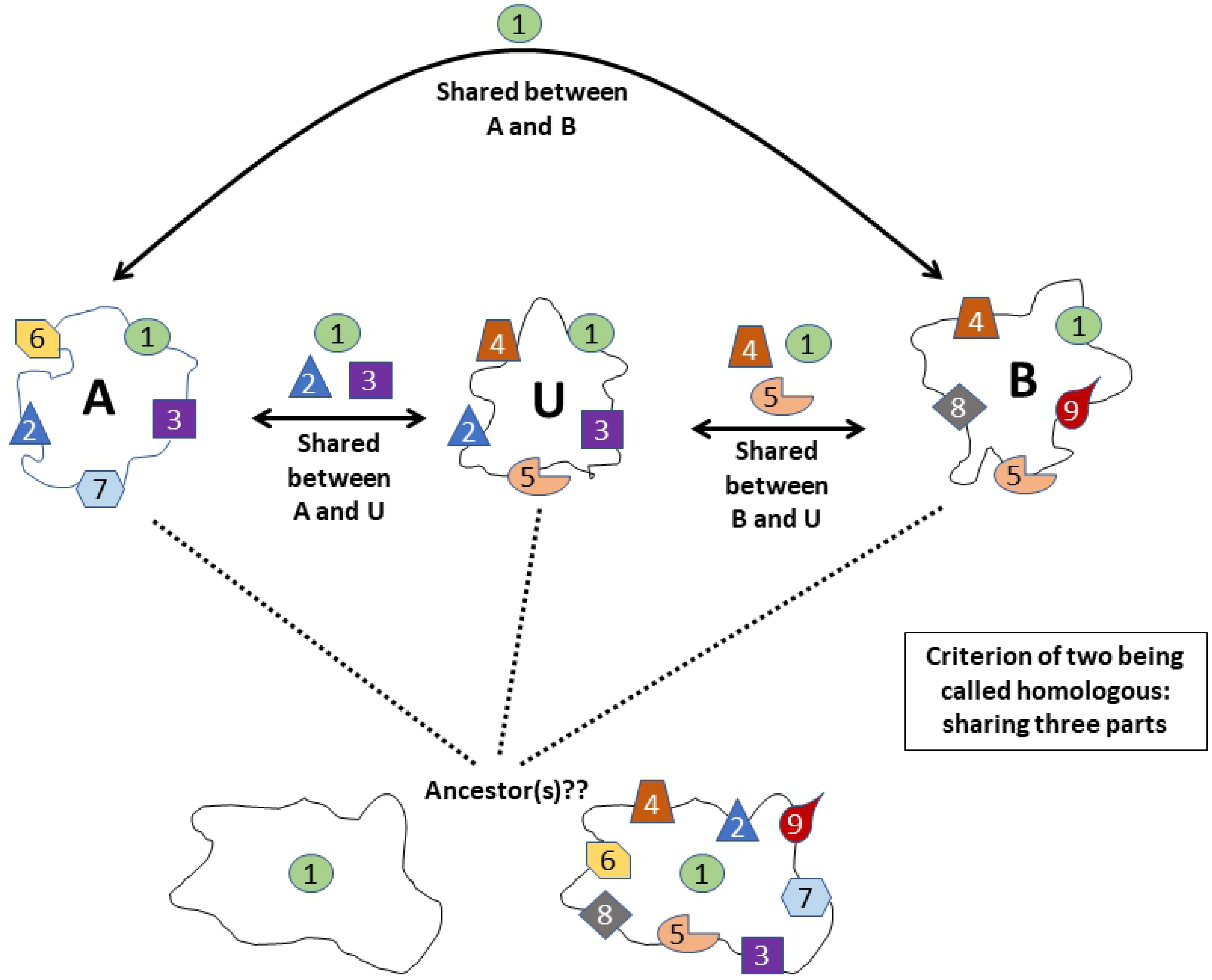
A possible scenario of how a homolog of a universal gene could be a non-universal gene.

### Different Levels of Homology

The puzzle that homologs of a universal gene are normally non-universal genes necessitates distinguishing different levels of homology. The concept of gene homology and the identification of homologous genes among different species are foundational to our study and numerous other comparative genomic studies. We think that it will be fruitful to define and distinguish different levels of homology. Here we propose the following levels of homology: 1) protein-domains (or partial sequence homology, or regional homology), 2) full length proteins, 3) full length genes, including the 5’- and 3’-untranslated regions and introns, 4) signal transduction pathways, 5) tissues, 6) organs, 7) body parts. The first three levels are at the molecular level and concern protein-coding genes. The higher the level of homology two genes share, the more likely they will perform the same function. Each level has its own value, even the lowest level, the level one. For instance, if two proteins both have a kinase domain, then they will be a kinase of some sort. Their other non-homologous regions may determine their substrates and their cellular locations of functioning. If these two proteins share homology throughout their entire length, then we can predict that they possess the same substrate specificity and function in the same cell compartment.

Currently, it is a general practice to call two proteins homologous as long as they share level one homology and be put into the same protein family. This can cause unnecessary challenges for functional annotation of genes because thus assigned family members may perform opposite or unrelated functions. This can be confusing, even misleading, especially when thus-identified “homologous” genes are given the same or similar names. For example, Frizzled, a seven-transmembrane protein, and FrzB (also known as soluble or secreted frizzled-related proteins), a protein that is similar to the amino-terminal cysteine rich domain of Frizzled but has no transmembrane segments, are included in the same protein family (InterPro: https://www.ebi.ac.uk/interpro/entry/InterPro/IPR015526/, PANTHER: http://www.pantherdb.org/panther/family.do?clsAccession=PTHR11309, UniProtKB: https://www.uniprot.org/uniprot/Q92765). However, the former is a Wnt receptor necessary for Wnt signaling, while the latter inhibits Wnt-signaling. A similar example is the *C. crescentus* peptide chain release factor 3 and the elongation factor G discussed earlier. They are put into the same subfamily of GTPases but perform different functions. Many bacterial and archaeal genes are called globins because they share partial sequence similarity with hemoglobins, but instead of carrying oxygen around like a hemoglobin, a bacterial globin does not bind oxygen, instead may function as a nitrogen monoxide detoxifier (39).

We propose to provide the following information when declaring two genes homologous: 1) level of homology, or homology coverage, 2) degree of homology, 3) e-value cutoff, and 4) a visual representation of the homology. Homology coverage should indicate whether the homology level is of protein-domain, full-length protein, or full-length DNA sequence. For example, we may divide every protein into four quarters, 1 to 4 from the N-terminal to the C-terminal. The homology between FrzB and Frizzled can be described as FrzB-p1-4/Frizzled-p1, “ p” for protein. Homology degree can be indicated with the percent identity of a BLASTp search. A visual representation of the homology should include both the regions that can be aligned and those that cannot be aligned. Fig 8 depicts a visual representation of homology between two subunits, Rpb A’ and A”, of archaea *Pyrococcus furiosus* RNA polymerase and the largest subunit Rpb1 of *S. cerevisiae* RNA polymerase II. A’ aligns with the first two quarters of Rpb1. A” aligns with the third quarter of Rpb1. The C-terminal quarter of Rpb1 is unique to eukaryotes and is essential for eukaryotic gene transcription initiation, elongation, termination and intron splicing (40). Specific information about the levels at which proteins are homologous will not only avoid making incorrect connection of protein functions but also facilitate understanding of true relationship of genes.

**Fig 8.**
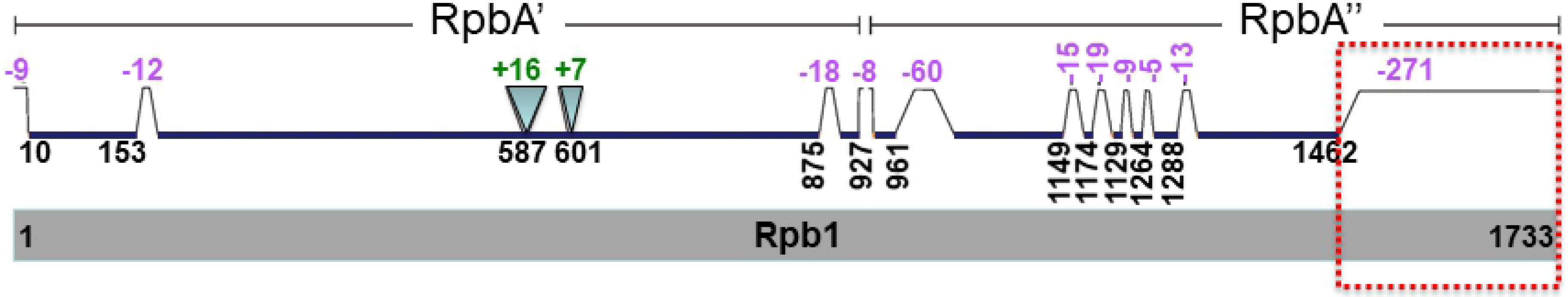
A comparison of S. cerevisiae Rpb1 (gray, bottom) with P. Furiosus Rpb A’ and A” (green, top). Segment locations are based on amino acid positions of Rpb1 protein. Segments present in Rpb1 but not in Rpb A’ and A” are indicated with trapezoids and negative numbers, while those segments absent in Rpb1 but present in Rpb A’ and A” are indicated with inverted triangles and positive numbers. The values of the numbers, which correlate with the sizes of the triangles and trapezoids, represent the numbers of amino acids of the corresponding regions that are present in only one of the two proteins compared. The C-terminal tail of Rpb1 that is missing in P. furiosus RNA polymerase is highlighted with a red dash-lined box. The comparison is from Fig 1I of (40).

### A Novel Index of Gene Diversity

Our birth-event-weighted gene distribution method can be used as a reasonable indication of the diversity of genes, or different types of genes. Since homologous genes (correctly defined as discussed above) tend to be similar, a non-discriminating counting would inflate the total gene number count encoded by all species on earth. The weighting approach corrects this, and thus can be used as an index of the number of gene types or gene diversity. Moreover, weighting is logically required if indeed homologous genes share a common ancestral gene. But this makes the explanation of the origin of ONO genes more acutely mysterious and makes it more important to study their functions, which we will address in the near future.

### Limitation of This Study

The number and identity of genes in each of our orphan, o+1, o+2… and universal homologous groups may be different using a different method and/or a different e-value cutoff for homology calling or when different species are included, though the trend of the differential growth of ONO and UNU genes will not change. It will be interesting to study how the phylogenetic profiles of proteins will change with a change of the criteria of homolog calling, such as the percentage of gene length covered, percentage of identity, alignment gap penalty, e-value cutoff, and calculation models (alignment algorisms) for these parameters. Furthermore, how the phylogenetic profiles would change when different species are included.

Recent years have witnessed a growth of interests in orphan genes (19–22, 28, 41–56). However, the increase of interests is incomparable to the increase of orphan gene number. Most of the orphans have unknown functions and will be a rich soil for discovery of new enzymes and/or unknown substrates of known enzymes (57) or new genotype-phenotype connections. The broad existence of orphan genes calls for a greater attention to them from the biological community.

## Conclusions

Our in-depth analysis of phylogenetic profiles of 317 proteomes across the three domains of life shows that ONO genes are a common occurrence in the sense that each organism has a significant number of them, while UNU genes are very rare. Most organisms, especially eukaryotes, do not have any UNU genes. Furthermore, the sum total of UNU genes almost plateaued while the number of ONO genes grew continuously when the number of organisms being analyzed increased. More importantly, every universal gene lost its status as a universal gene when the sampled number of organisms is increased. These results revealed a great challenge to explain not only the origin of genes but also the origin of life and the origin of biodiversity. We propose to use the birth-event-weighted distribution of genes as an indication of gene diversity, even though the weighting makes it more difficult to explain the origin of genes by the common belief that genes were generated via duplication and diversification because of the greatly enlarged portion of the ONO genes.

## Acknowledgments

The author thanks Andrew Jones for help with calculating the numbers of homologs, Rob Sadler and Paul Nelson for comments on the manuscript, Alan Marshall and Nicholas Valentine for help with computer.

## Additional information

**S1 Fig. Change of numbers of genes in different gene groups in 12 additional species**

Note that the first vertical line in each panel is not its Y-axis but the orphan genes in the corresponding organism. The numbers in the parenthesis next to the name of an organism is the number of its orphan gene, the total protein-coding gene in its genome, and the percent of the orphan genes of the total.

**S2 Table. Number of genes in different gene groups for all NMS**

**S3 Table. Distribution of orphan, nearly-orphan, universal, and nearly-universal genes of all NMS**. The 17 chosen species were highlighted with red font.

**S4 Table. Weighted distribution of orphan, nearly-orphan, universal, and nearly-universal genes of all NMS**

**S5 Table. Analysis of universal genes of selected organisms**. Reference genes are highlighted with red font. The seemingly out-of-place homologs are shaded with yellow.

**S6 Table. Analysis of universal genes of unknown identity**. Reference genes are highlighted with red font.

**S7 Table. Analysis of the identity of universal genes of all NMS**

